# Centromere Innovations Within a Mouse Species

**DOI:** 10.1101/2023.05.11.540353

**Authors:** Craig W. Gambogi, Nootan Pandey, Jennine M. Dawicki-McKenna, Uma P. Arora, Mikhail A. Liskovykh, Jun Ma, Piero Lamelza, Vladimir Larionov, Michael A. Lampson, Glennis A. Logsdon, Beth L. Dumont, Ben E. Black

## Abstract

Mammalian centromeres direct faithful genetic inheritance and are typically characterized by regions of highly repetitive and rapidly evolving DNA. We focused on a mouse species, *Mus pahari,* that we found has evolved to house centromere-specifying CENP-A nucleosomes at the nexus of a satellite repeat that we identified and term π-satellite (π-sat), a small number of recruitment sites for CENP-B, and short stretches of perfect telomere repeats. One *M. pahari* chromosome, however, houses a radically divergent centromere harboring ∼6 Mbp of a homogenized π-sat-related repeat, π-sat^B^, that contains >20,000 functional CENP-B boxes. There, CENP-B abundance drives accumulation of microtubule-binding components of the kinetochore, as well as a microtubule-destabilizing kinesin of the inner centromere. The balance of pro and anti-microtubule-binding by the new centromere permits it to segregate during cell division with high fidelity alongside the older ones whose sequence creates a markedly different molecular composition.

**Teaser:** Chromatin and kinetochore alterations arise in response to evolutionarily rapid changes to underlying repetitive centromere DNA.

## Introduction

Centromeres are the loci that coordinate chromosome segregation during cell division (*1*). They do so by assembling a proteinaceous structure, the kinetochore, at cell division that attaches to spindle microtubules, housing the chromatin that regulates microtubule attachment to ensure error-free segregation, and serving as the final site of sister chromatid cohesion. In many species, including mammals, the site for all of these functions is epigenetically specified by the presence of nucleosomes harboring the histone H3 variant, CENP-A.

Despite generally shared and essential functional roles there is marked diversity in the DNA sequences and molecular composition of centromeres between different eukaryotic species. Centromere formation can influence evolution by allowing some centromeres to be preferentially inherited during female meiosis by biasing segregation outcomes in a process called ‘centromere drive’ or more generally ‘meiotic drive’ (*2, 3*). Centromeres that direct biased segregation to the egg are referred to as ‘stronger’ centromeres. Among other factors, expanding the region of DNA housing CENP-A nucleosomes can strengthen the centromere (*4*). Female meiotic drive is thought to be the major driver of rapid evolution of centromeric DNA (*5*).

One powerful model system for assessing the molecular basis for female meiotic drive is the mouse (*6*). Prior work has demonstrated that major differences in the abundance of repetitive centromere DNA between inbred laboratory strains or species lead to differences in which chromosomes are more likely to be inherited through meiosis (*6*). While centromere DNA sequence and architecture differs between mouse species, in each of the reported cases, centromere DNA differences between chromosomes within a strain or species are thought to be negligible (e.g. every *Mus spretus* chromosome has a nearly identical repeat at each centromere at a similar abundance) (*4, 7, 8*). Of course, centromeres are present on separate chromosomes implying that DNA sequence-based differences between centromeres are homogenized across the genome through some undefined selective pressure to do so. More precisely, individual chromosomes are physically unlinked and subject to the independent accrual of new mutations. Non-allelic homologous repair processes can homogenize centromeres from different chromosomes, erasing signals of chromosome-level centromere divergence (*9, 10*). Such mechanisms have likely been particularly active on acrocentric mouse chromosome, where centromeres colocalize at the nuclear periphery during meiosis onset, prior to the completion of double-strand DNA break repair (*11*). Nonetheless, the rapid evolution of centromeric DNA suggests that genomes with heterologous centromere composition are potentially pervasive, even if only transiently manifest in mouse genomes.

Thus, evolutionary intermediates must have existed prior to homogenization, and the molecular consequences remain unclear of having divergent centromeric DNA within a single mouse strain and/or species. In other eukaryotes, there are examples of different centromeres within a species, but it is unclear how they relate to the mouse model for strengthening through modulation of DNA repeat number or sequence due to major differences at the centromere. For instance, plant neocentromeres, like a famous example in maize (*12–14*) can function not through an actual centromere/kinetochore but by directing independent movements through tethering a specialized motor protein to the spindle. In mammals, evolutionarily young centromeres have been found on up to half of the centromeres of individual equine species (*15*). Further, many are present (albeit on smaller numbers of chromosomes) in several other vertebrate systems (*16–19*). In all these documented cases, the young centromeres consist of non-repetitive DNA. Given the recent successful studies using the mouse model system to reveal the role of centromere strength in centromere evolution (*4, 8, 20*), advances in mice on isolating and studying new radical changes in repetitive centromere DNA are likely to have important implications for advancing models of centromere evolution in diverse eukaryotic species.

CENP-B is the only known sequence-specific DNA-binding protein found at many eukaryotic centromeres, including at the centromeres in diverse mammalian species. It recognizes a conserved 17-mer sequence termed the CENP-B box, in which 9 positions are essential for CENP-B binding (*21, 22*). The CENP-B box is found within the sequences of the centromere repeat monomers (i.e. within the 171 bp alpha-satellite repeat in *Homo sapiens* and within the 120 bp repeat in minor satellite in *Mus. musculus*) (*23, 24*). While not essential for centromere function (indeed CENP-B boxes are absent on the Y chromosome in humans and mice (*25*)), CENP-B can buffer against other molecular insults and is a prime candidate to play a role in modulating centromere strength (*20, 25, 26*). CENP-B serves to support the pericentromeric enrichment of constitutive heterochromatin (i.e. chromatin enriched with nucleosomes marked with histone H3 lysine 9 trimethylation [H3K9me3]) that, in turn, enhances the recruitment of inner centromere components involved in sister chromatid cohesion and the process of mitotic error correction (*20, 27–29*). CENP-B, likewise, enhances kinetochore formation through its ability to bind an essential centromere protein, CENP-C (*25*). Removal of CENP-B enhances functional differences in female meiosis between diverged strains of *M. musculus* that have approximately ten-fold differences in minor satellite abundance relative to one another (*20*). Thus, there is a strong support for the notion that CENP-B can play a key role in modulating centromere strength.

Here, we find that CENP-B is dispensable for CENP-A nucleosome positioning on minor satellite DNA, suggesting that its roles are likely limited to strengthening the centromere by other proposed means that rely on the amount of CENP-B at the centromere. We then identify a single chromosome in the mouse species *M. pahari* that has a massive expansion of a newly evolved repeat array that houses >20,000 functional CENP-B boxes: ∼100-fold more than on the other *M. pahari* centromeres. Using a comprehensive set of short and long-read sequencing-based methodologies, we define this centromere and the more typical centromeres in *M. pahari*. The latter accumulate kinetochore forming CENP-A chromatin at a subset of repeats that harbor a relatively small number (hundreds) of CENP-B boxes, as well as up to 68,000 telomere repeats. Taken together, our sequencing efforts predict a difference in the molecular composition of the two types of centromeres within a single organismal genome. We test this notion and determine how the opposing recruitment of microtubule-binding and microtubule-destabilizing factors co exist in the same mouse species.

## Results

### Positioning of CENP-A Nucleosomes on Minor Satellite is Independent of CENP-B

In *M. musculus*, CENP-B boxes are only found within minor satellite DNA (*30*). CENP-A nucleosomes predominantly occupy a single site within the minor satellite repeat, with their centers (also known as the nucleosomal dyad) precisely positioned within the CENP-B box (*4*). While this nucleosome position is used by conventional nucleosomes containing canonical histone H3, it is only one of several prominent sites for CENP-A nucleosome assembly. In principle, minor satellite DNA sequence could directly impact the position of CENP-A containing nucleosomes independently of CENP-B binding. Alternatively, the CENP-B protein could impact CENP-A nucleosome positioning upon binding to the CENP-B box and through its direct and indirect interactions with the CENP-A nucleosomes. To distinguish between these possibilities, we enriched for nucleosomes containing either CENP-A or H3K9me3 via chromatin immunoprecipitation (ChIP) from chromatin isolated from wild type (WT; C57BL/6J) or CENP-B^-/-^ (C57BL/6J) mice. We found that positioning on minor satellite of CENP-A nucleosomes, H3K9me3 nucleosomes, and the total pool of nucleosomes (input to the native ChIP) were essentially unchanged in the absence of CENP-B protein (Fig. 1A, B). Thus, our data support the notion that minor satellite DNA sequence is uniquely responsible for positioning of CENP-A nucleosomes, independently of the presence of CENP-B protein. Our results suggest that CENP-B protein, the CENP-B box, and centromere satellite sequences are important for us to consider in contributing to centromere drive.

**Figure 1:**
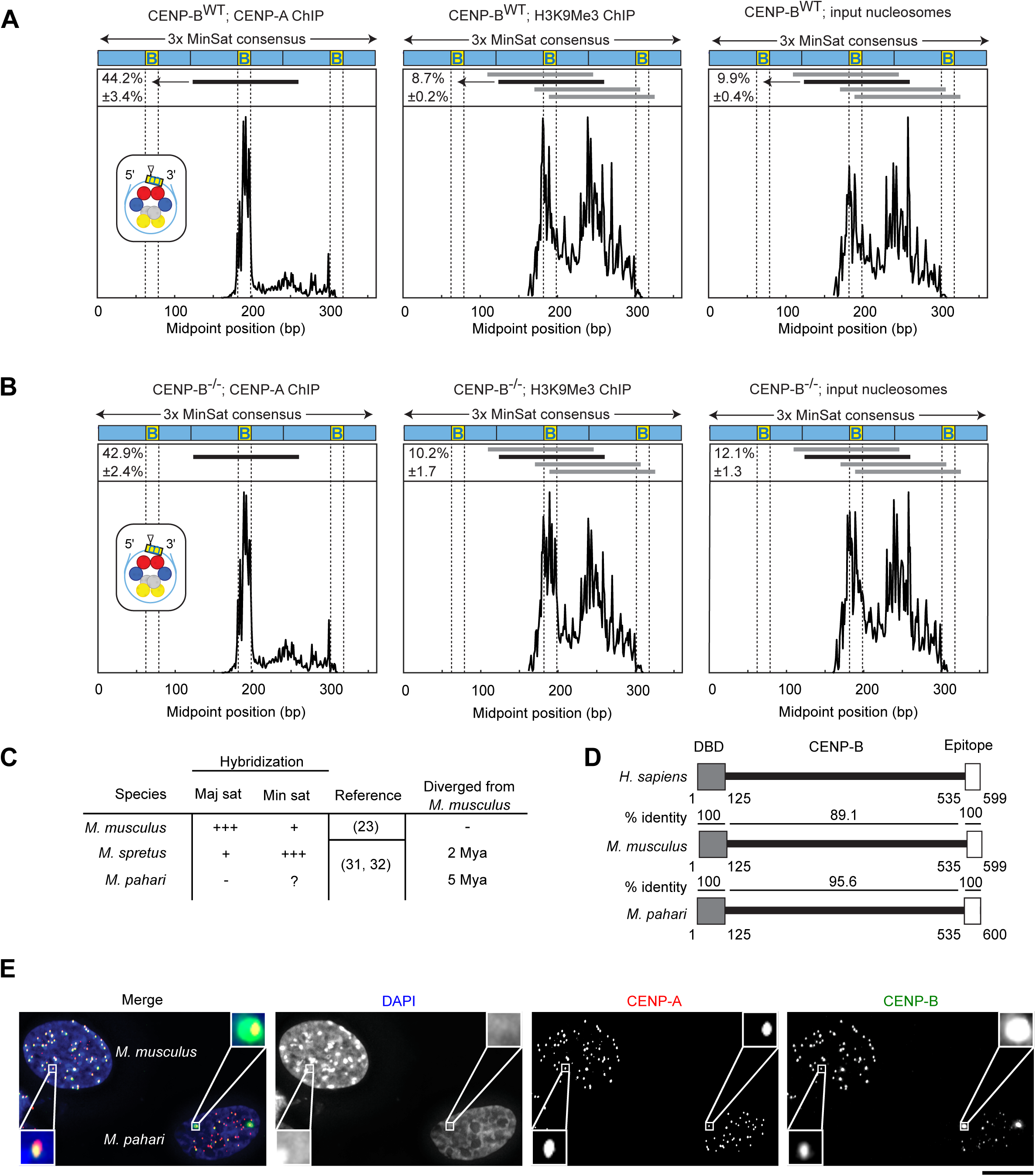
CENP-B occupancy on centromere DNA does not impact CENP-A nucleosome phasing, but does vary widely between and within mouse species. A) Midpoint position of CENP-A ChIP H3K9Me3 or input reads (size 100–160 bp) from WT *M. musculus* along the trimer minor satellite consensus sequence. Vertical lines indicate the 17-bp CENP-B box. The major CENP-A nucleosome position (identified in the CENP-A ChIP samples) is indicated by a horizontal black line above the respective midpoint values and schematized (inset) for CENP-A ChIP with a triangle representing the dyad position. The same nucleosome position is indicated in the H3K9Me3 and input samples. Numbers to the left of the positions indicate the percentage of reads (mean ± SEM; n = 3 independent experiments) where the midpoint spans the 10 bp at the 3’ end of the CENP-B box (yellow, labeled B). Horizontal gray lines indicate other major nucleosome positions in the H3K9Me3 and input samples. B) Midpoint position of CENP-A ChIP H3K9Me3 or input reads (size 100–160 bp) from CENP-B KO *M. musculus* along the trimer minor satellite consensus sequence. C) Centromere satellites from *M. musculus, M. spretus*, and *M. pahari*. D) CENP-B is highly conserved in mouse species, with 100% identical sequences in both the DNA binding domain and the epitope targeted by the CENP-B antibody used in our study. E) Immunofluorescence of CENP-A and CENP-B from lung fibroblast cells derived from *M. musculus* (with their nuclei identified by strong DAPI-staining pericentromeres) or *M. pahari*. Bar, 10 μm.

### Rapid Centromere DNA Repeat Evolution Impacts the Amount of CENP-B at Centromeres

Early hybridization studies indicate that closely related house mouse species are undergoing evolutionarily rapid changes in centromere DNA sequence (Fig. 1C) (*23, 31, 32*). This divergence can include the number of CENP-B boxes and/or the sequence of the repeat, itself (*33*). One way to alter CENP-B box number is to vary the abundance of homogeneous centromere repeats. For instance, in *M. spretus* minor satellite is the most abundant centromere satellite and major satellite is much less abundant (*23*); the opposite of what is found in *M. musculus*. Changes also include apparent drastic alterations in DNA sequence, as in *M. pahari* where major satellite is undetectable (*31, 32*).

Two initial observations suggested that investigating the centromere diversity in *M. pahari* could yield new insights into the mechanism governing centromere strength. First, the *M. pahari* genome encodes the CENP-B protein which is almost identical to its counterpart in *M. musculus* and 100% identical in its DNA-binding domain (Fig. 1D). Such high species-level protein conservation is highly unlikely to persist over evolutionary time in the absence of purifying selection to retain CENP-B function. Thus, we anticipated that *M. pahari* centromeres would contain repeats—minor satellite DNA or other divergent repetitive centromere DNA—that harbor functional CENP-B box sequences capable of CENP-B binding. Second, we found that while most *M. pahari* centromeres have low (relative to those from *M. musculus* cells co-seeded for immunofluorescence measurements) yet detectable levels of CENP-B, a pair of very strong foci of CENP-B are present (Fig. 1E). We concluded that the pair of foci likely represent a single pair of homologous chromosomes. Thus, our initial observations suggested that in *M. pahari*, major changes exist in centromere DNA both relative to *M. musculus* and between different *M. pahari* chromosomes and that impacts the CENP-B abundance at the centromeres.

### Identification of a Divergent Centromere Satellite, π-sat

Since no centromere satellite has been identified in *M. pahari*, we employed several strategies to identify candidate centromere repeats (Fig. 2A). The first strategy was a *k*-mer-based approach using an existing short-read sequencing data set (*34*). This yielded a top hit with a repeat unit length of 189 bp (Fig. S1). The second strategy was an analysis of total nucleosomal DNA and CENP-A nucleosome-enriched (native ChIP) short-read data with the computational pipeline TAREAN (*35*) coupled to downstream analysis of native Oxford Nanopore Technologies (ONT) long-read sequencing we performed of the *M. pahari* genome (Fig. 2A; see Methods for details of the strategy we employed). This produced a total of three sequences with a high likelihood of satellite DNA (Fig. 2B; Methods). Of the three sequences, the 189 bp satellite, which we term π-sat, is nearly identical to the top hit identified by the *k*-mer strategy (Fig. S1). Consistent with our hypothesis that is a centromere repeat, π-sat hybridizes to a single locus on each chromosome in a chromosome spread of mitotic *M. pahari* cells (Fig. 2C). However, the π-sat sequence lacks an intact CENP-B box (Fig 2D). The two remaining repeats we identified are related to π-sat: one (π-sat^sh^) is ∼50 bp shorter, whereas the other (π-sat^B^) contains an intact CENP-B box (Fig. 2D).

**Figure 2:**
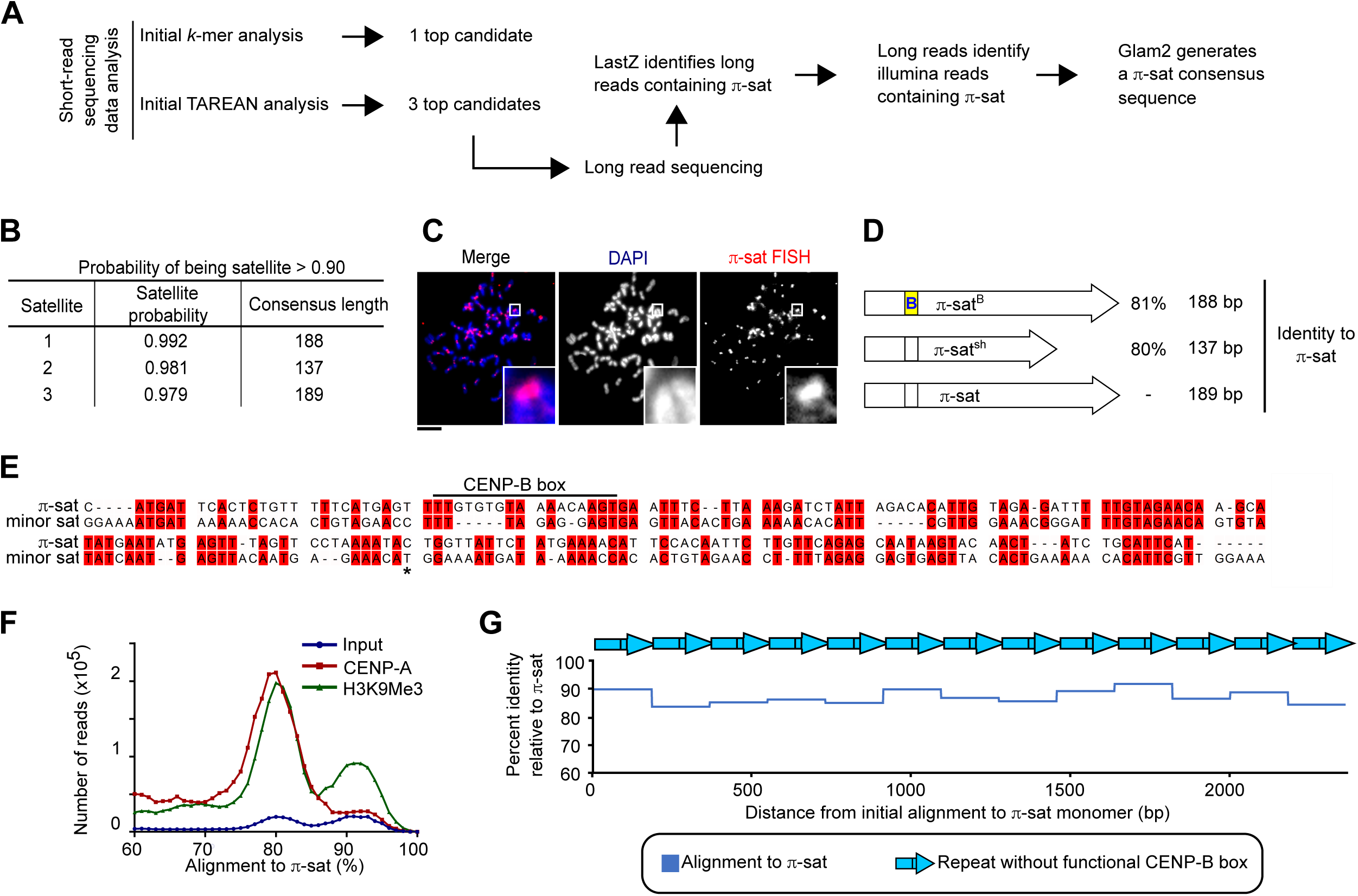
Identification of the most abundant form of centromere repeats *in M. pahari*: π-sat. A) Two approaches to identify *M. pahari* centromeric repeats. B) Satellite sequences derived from TAREAN analysis on input sequencing data. Satellite probability was calculated as described in Methods. C) Representative image of *M. pahari* centromeric DNA labeled with FISH probe using consensus sequence derived from *k*-mer approach. Insets: 7.9x magnification, Bar, 10 μm. D) Schematized representation of the 3 satellites identified by TAREAN analysis. E) Alignment of the π-sat consensus sequence to minor sat consensus sequence. A dimer of π-sat was aligned to a trimer of minor satellite and the first monomer of π-sat is shown. The end of first monomer of minor satellite is marked with an asterisk. F) Histograms show distribution of reads from input, CENP-A ChIP, or H3K9Me3 ChIP aligning to π-sat. G) Representative example of a π-sat containing ONT long read that was divided into monomers. The percent identity of each monomer to π-sat is plotted.

We noted that none of the three π-satellites we identified were closely related to major satellite from *M. musculus*, explaining why early hybridization studies failed to identify major satellite in *M. pahari* (*31, 32*). Minor satellite similarly has only small regions of identity with π-sat, and the region of π-sat aligning to the CENP-B box has several substitutions (Fig. 2E). Alignment of enriched sequences from CENP-A (functional centromere) and H3K9me3 (enriched in pericentromeric heterochromatin) native ChIP with π-sat yielded strong peaks of high sequence identity (Fig. 2F). Further, we noted that many of the long-reads that align to π-sat consisted of homogenous stretches where π-sat contained no intervening sequences, including any π-sat^sh^ or π-sat^B^ (Fig. 2G). Together with the FISH and native ChIP data, these experiments suggest that most or all *M. pahari* centromeres harbor long and uninterrupted stretches of π-sat repeats that lack functional CENP-B boxes.

### A Chromosome Pair with Highly Homogenized π-sat^B^

To gain an understanding of the centromere sequences that harbor functional CENP-B boxes, we employed another strategy (Fig. 3A), starting with native ONT long reads that harbor functional CENP-B box sequences. This approach yielded a refined centromere consensus sequence (Fig. 3B) that corresponded to what we had initially identified as π-sat^B^ (Fig. 2). CENP-A and H3K9me3 native ChIP reads contained many sequences that align well to the π-sat^B^ consensus sequence (Fig. 3C). Peaks around 83-86% sequence identity likely correspond to alignments with general π-sat, while a peak around 94-96% sequence identity likely represents π-sat^B^ sequences. We designed a FISH probe using the π-sat^B^ consensus sequence and found that it hybridized to a pair of mitotic chromosomes in *M. pahari* cells (Fig. 3D). Further, in interphase *M. pahari* cells the π-sat^B^ probe colocalized with a probe specific to the CENP-B box (Fig. S2). This supported our prior conclusion that the two nuclear puncta with high amounts of CENP-B (Fig. 1E) correspond to a single pair of homologous chromosomes. Alignment of sequences found on long reads containing either π-sat or π-sat^B^ showed that π-sat^B^ has near invariance at the CENP-B box positions that are required for CENP-B binding, including at the positions that diverge from the π-sat consensus (Fig. 3E). The majority of ONT long reads containing centromere repeats were homogenous stretches that align more closely to π-sat and were devoid of CENP-B boxes. On the other hand, a smaller proportion contain centromere repeats that, while also comprising homogenous stretches, contain many functional CENP-B boxes and align more closely to π-sat^B^ (Fig. 3F). Our findings indicate that a homologous pair of chromosomes that bind high levels of CENP-B harbor a large and highly homogenous derivative of the satellite present on the other chromosomes.

**Figure 3:**
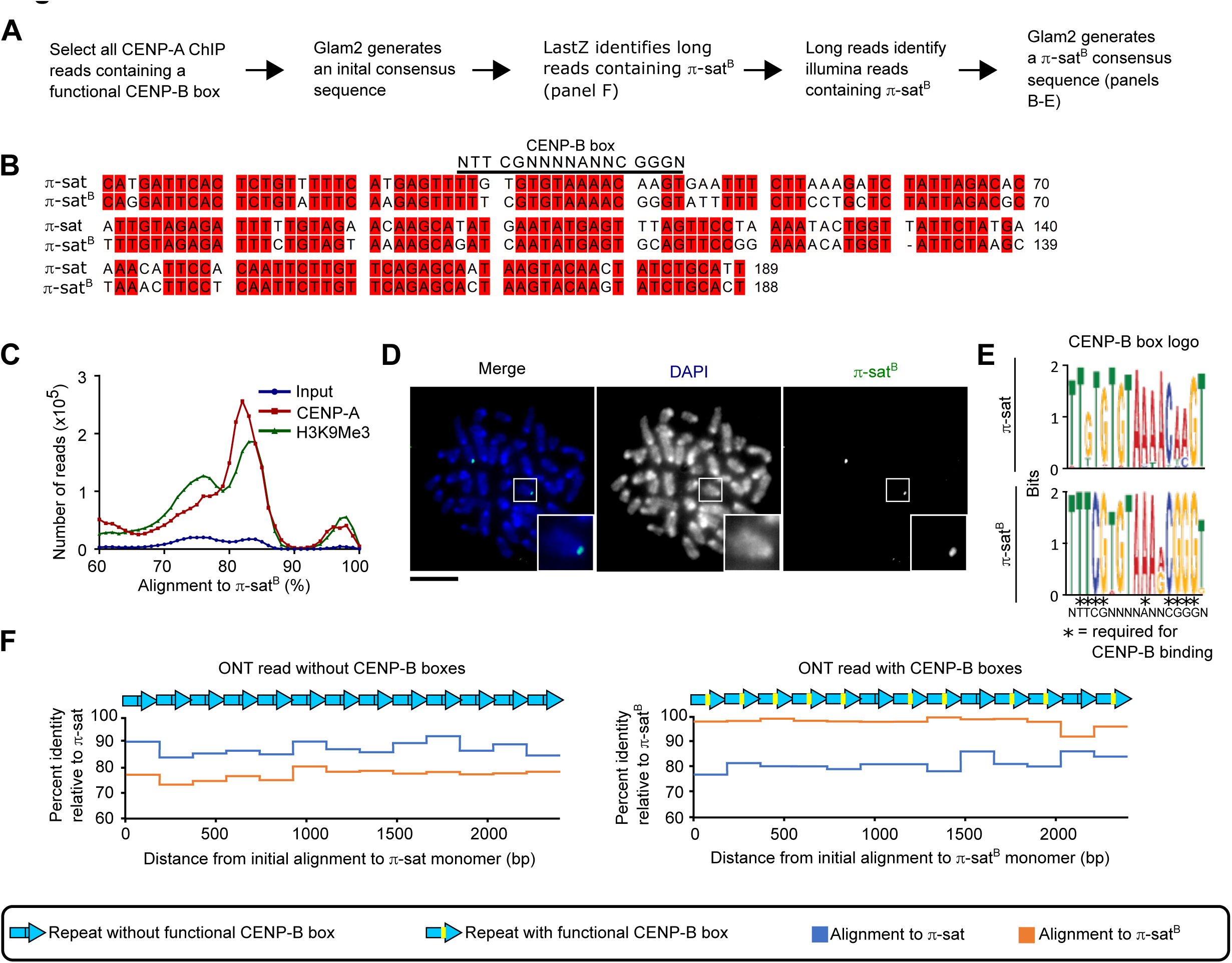
π-sat^B^ is highly homogenous, restricted to a single pair of chromosomes, and present in long, contiguous blocks that lack generic π-sat. A) Approach to identify CENP-B box containing satellite. B) Alignment of π-sat and π-sat^B^. C) Histograms show distribution of reads from input, CENP-A ChIP or H3K9Me3 ChIP aligning to π-sat^B^ . D) Representative image of *M. pahari* centromeric DNA labeled with FISH probe using π-sat^B^ consensus sequence. Insets: 2.5x magnification, Bar, 10 μm. E) Logo representation of the CENP-B box consensus of π-sat and π-sat^B^. F) Plots of the percent identity of satellites along a portion of representative ONT reads with (right) and without (left) CENP-B boxes to the π -sat and π -sat^B^ consensus sequences.

### High-accuracy Sequence Assemblies of M. pahari Centromeres

In order to identify the chromosome with high amounts of CENP-B, as well as to more broadly understand centromere structure in *M. pahari*, we set out to generate centromere sequence assemblies from several *M. pahari* chromosomes. While murine centromeres have long been assumed to be relatively intractable to sequence assembly due to high repeat homogeneity and apparent lack of higher-order repeat patterns (e.g., this is true of the best known murine centromere repeat for centromere function in cell division, minor satellite from *M. musculus*), we were encouraged by two aspects. The first was the success of Pacific Biosciences high-fidelity (Pacbio HiFi) long-read sequencing in assembling human centromeres with high accuracy (*36*– *38*). The second was our finding that π-sat is not as homogenous as minor satellite (Fig. 2G). Our initial focus for sequence assembly was of the chromosome containing a large array of π-sat^B^ (Fig. 4A). Therefore, we generated a 22-fold coverage of PacBio HiFi data from the *M. pahari* genome and assembled it with the whole-genome assembler hifiasm (*39*). This generated a whole sequence assembly that was 4.54 Gbp in length, consistent with its diploid nature, and containing a contiguous assembly from the telomere through the first 13 Mbp of the chromosome containing arrays of π-sat^B^. Aligning this contig to an adjacent contig was sufficient to extend to complex sequence that matches chromosome 11 from the initial genome build of *M. pahari* (*34*). This chromosome is telocentric, with no intervening sequence between perfect telomere repeats and centromeric repeats (Fig. 4A).

**Figure 4:**
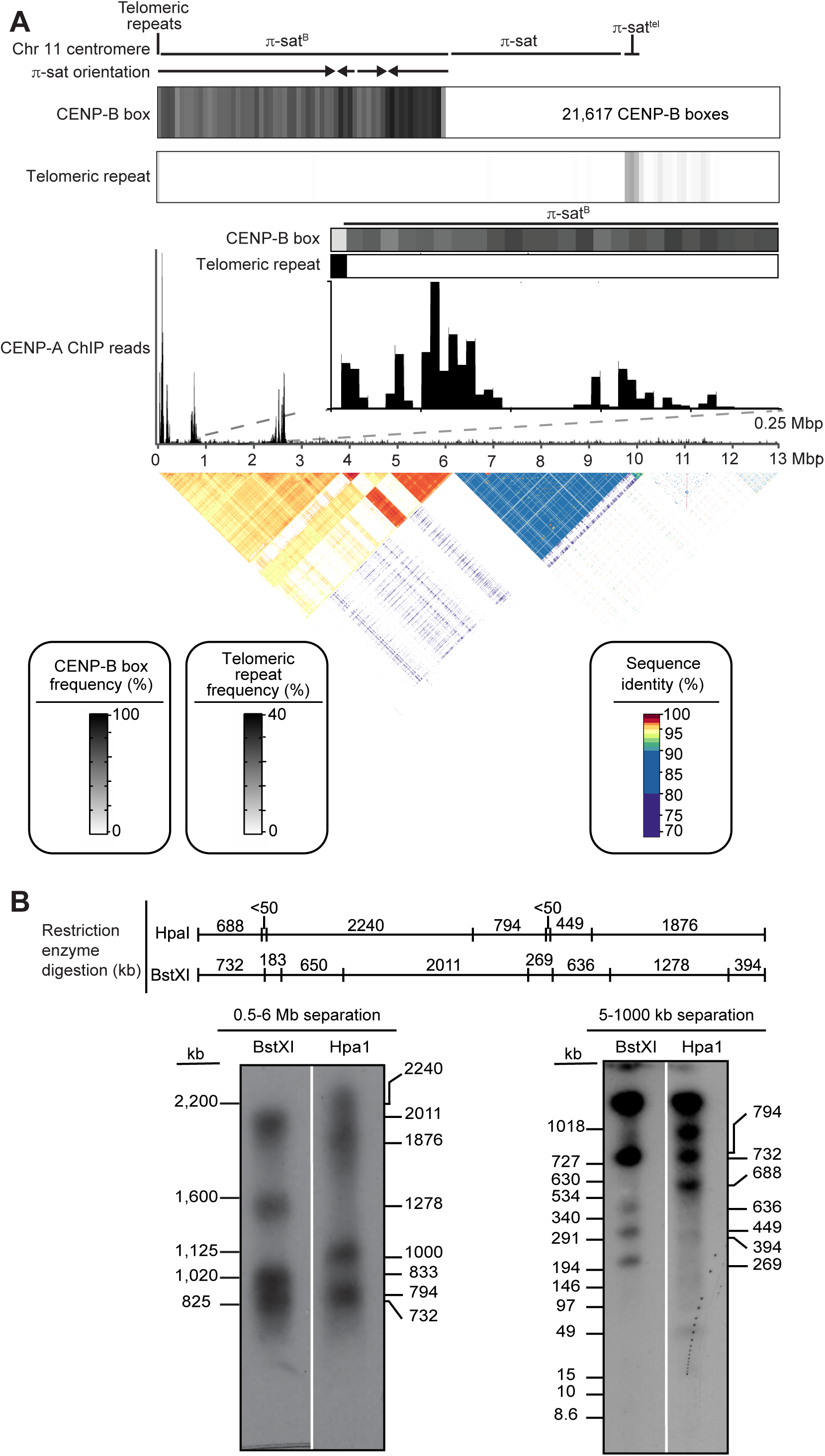
Genomic assembly reveals the identity and nature of the centromere harboring π-sat^B^. A) The composition of the centromere of chromosome 11. The assembly consists of, in order, 8 kb of telomeric repeats, 6 Mbp of π-sat^B^, 3.6 Mb of π-sat, 400 kb of π-sat^tel^, followed by other repetitive elements. The total number of CENP-B boxes (21,617) on this centromere is denoted. The fraction of π-sat repeats containing a functional CENP-B box (NTTCGNNNNANNCGGGN) and the frequency of telomeric repeats (TTAGGG) are shown. CENP-A ChIP-seq reads were aligned to the chromosome 11 centromere assembly. A pairwise sequence identity heat map indicates that the centromere consists of 6 Mbp of highly homogenous π-sat^B^. B) Schematic of predicted restriction digest sites of chromosome 11 with BstXI and HpaI. Pulsed-field gel Southern blot of *M. pahari* DNA confirms the structure and organization of the chromosome 11 centromeric HOR array. For each gel, left corresponds to ethidium bromide (EtBr) staining and right, ^32^P-labelled chromosome 11 p-sat^B^ specific probe. The left gel was run at conditions to separate DNA from 0.6-5 Mb and the right gel was run at conditions to separate DNA from 5-1000 kb.

The first centromere repeats consist of a ∼6 Mbp block of contiguous π-sat^B^. The first 3.7 Mbp of this π-sat^B^ array includes monomers in exclusive head-to-tail orientation. The directionality of the head-to-tail repeats switches three times over the next 2.3 Mbp. In total, this 6 Mbp block houses 21,617 functional CENP-B boxes (Fig. 4A) explaining the massive enrichment of CENP-B on this chromosome (Fig. 1E). A ∼4 Mbp contiguous stretch of π-sat lies distal to the π-sat^B^ arrays, followed by a shorter stretch of π-sat variant, π-sat^tel^ (Fig. 5F). π-sat^tel^ is a more complex composite repeat monomer comprised of elements built from π-sat, π-sat^sh^, and 2-16 telomere repeats. CENP-A association is not uniform across the chromosome 11 centromere, with enrichment localized to three sites: a site of enrichment adjacent to the telomere (0-250 kbp) and two more regions marked by peaks at ∼750 kbp and 2.5 Mbp from the telomere, respectively (Fig. 4A). CENP-A peaks are only observed on π-sat^B^, but not on π-sat or π-sat^tel^ (Fig. 4A). Southern blots of *M. pahari* DNA digested with two restriction enzymes, BstXI and HpaI, and probed with π-sat^B^ almost perfectly match the pattern predicted by our assembly (Fig. 4B). Two predicted bands (183 and 650 kb) for BxtXI digestion were not detected, but one at 833 kb was (Fig. 4B). This minor difference is likely due to a sequence polymorphism between the animal used to generate the assembly versus the one used to harvest DNA for the blot. Thus, despite the high degree of sequence identity between repeat monomers and the lack of other unique sequences for a >6 Mbp span of π-sat^B^, our approach with PacBio HiFi long read sequencing downstream assembly strategy is extremely faithful.

**Figure 5:**
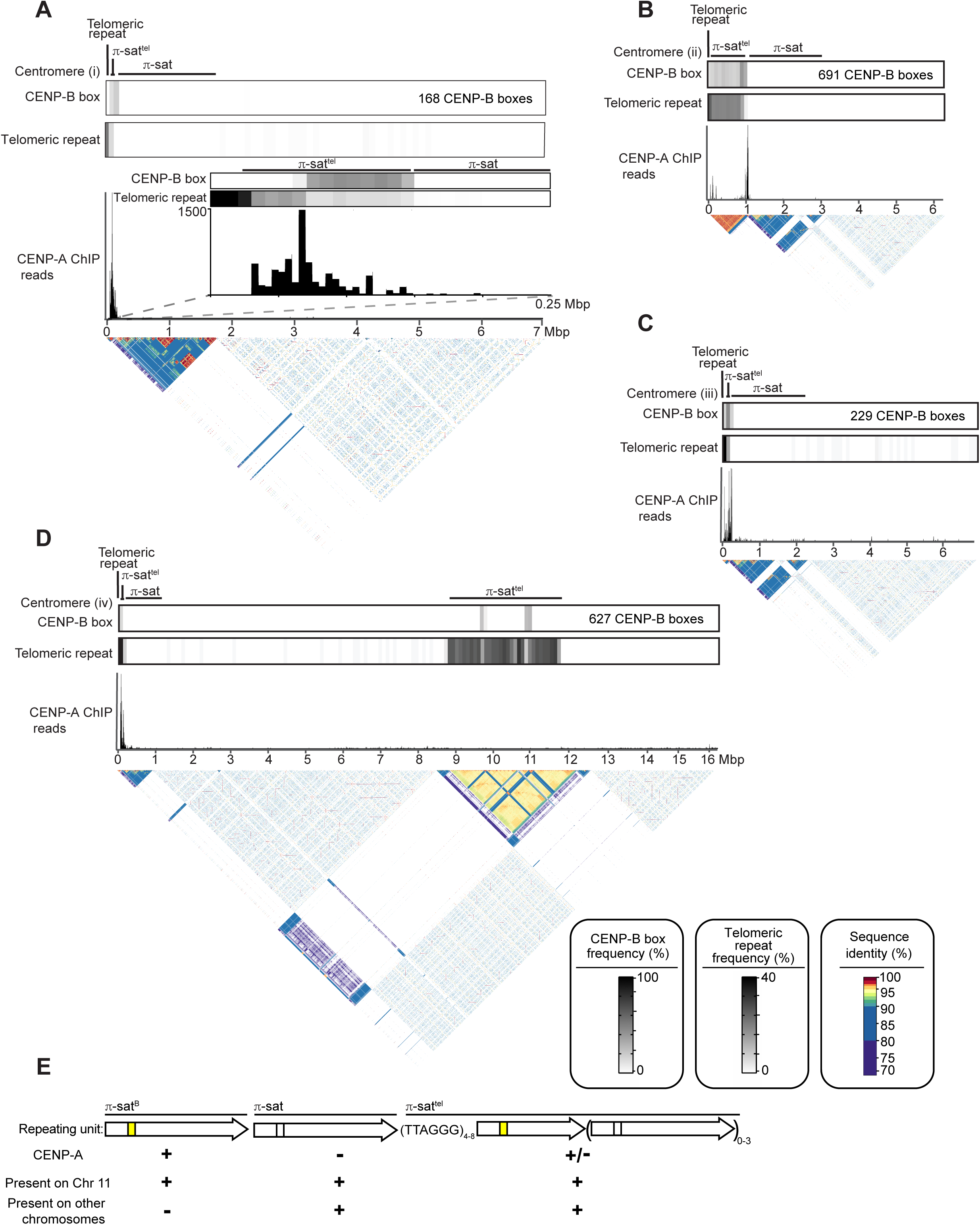
Evolutionarily older *M. pahari* centromeres harbor CENP-A nucleosomes near CENP-B boxes and π-sat^tel^. A-D) The composition of a representative *M. pahari* centromeres. Each of the assembly consists of in order an array of telomeric repeats, an array of π-sat^tel^, and an array of π-sat followed by various repetitive elements. The fraction of π-sat repeats containing a functional CENP-B box (NTTCGNNNNANNCGGGN) and the frequency of telomeric repeats (TTAGGG) are shown. CENP A ChIP-seq reads were aligned to the assembly revealing that CENP-A is primarily present on π-sat^tel^. A pairwise sequence identity heat map indicates the degree of homogeneity in centromeric DNA. E) The types of repeating units found at *M. pahari* centromeres.

We successfully assembled seven other *M. pahari* centromeres (Fig. 5A-D, S4). Note, all seven have unmapped regions between the centromere and the rest of the chromosome that preclude assignment to a particular *M. pahari* chromosome, so we have numbered them centromere (i)-(vii). They vary in size and precise arrangement, are commonly telocentric, and house π-sat^tel^ between the telomere and a long stretch of π-sat (Fig. 5A-D, S4). Importantly, none contain π-sat^B^ (Fig. 5A-D, S4). CENP-A peaks are almost entirely restricted to π-sat^tel^, as are functional CENP-B boxes (Fig. 5A-D, S4). The functional CENP-B boxes are almost exclusively confined to π-sat^tel^ repeats and vary in their sequence from those found on chromosome 11 in π-sat^B^ and are much less abundant (Fig. 5A-D, S3, S4). The majority of the π-sat repeats harbor non-functional CENP-B boxes that do not match the consensus required for CENP-B binding (Fig. 5A-D, S4). Thus, other assembled centromeres harbor 27-143 times fewer total functional CENP-B boxes than chromosome 11. For all centromeres that we assembled, the major site of CENP-A enrichment spans 100-300 kbp (Fig. 5A-D, S4). As far as the role of the different specific forms of π-sat, general π-sat is the most abundant and represent a candidate pericentromeric satellite (analogous to major satellite DNA in *M. musculus*), while both π-sat^tel^ and π-sat^B^ are primary sites for kinetochore forming chromatin containing CENP-A nucleosomes (Fig. 5E). Compared to chromosome 11, the other centromeres contain π-sat wherein monomer units are less similar to one another (Fig. 3F). Thus, it appears that the highly homogenous chromosome 11 centromere is evolutionarily more recent. In total, our long-read analysis of *M. pahari* centromeres define the general sequence features of *M. pahari* centromeres, including the evolutionary young centromere on chromosome 11.

### Co-existence of Chromosomes with Markedly Different Abundance of Centromere Factors

Chromosome 11 has a markedly different centromere repeat that leads to massive differences in CENP-B abundance (Fig. 4A). To test whether or not the large difference of CENP-B leads to higher levels of H3K9me3 accumulation, we performed quantitative immunofluorescence on interphase cells (Fig. 6A,B). Indeed, chromosome 11 has 1.6-fold higher H3K9me3 relative to that measured at the centromeres of other chromosomes have low yet detecteable CENP-B levels (Fig. 6A,B).

**Figure 6:**
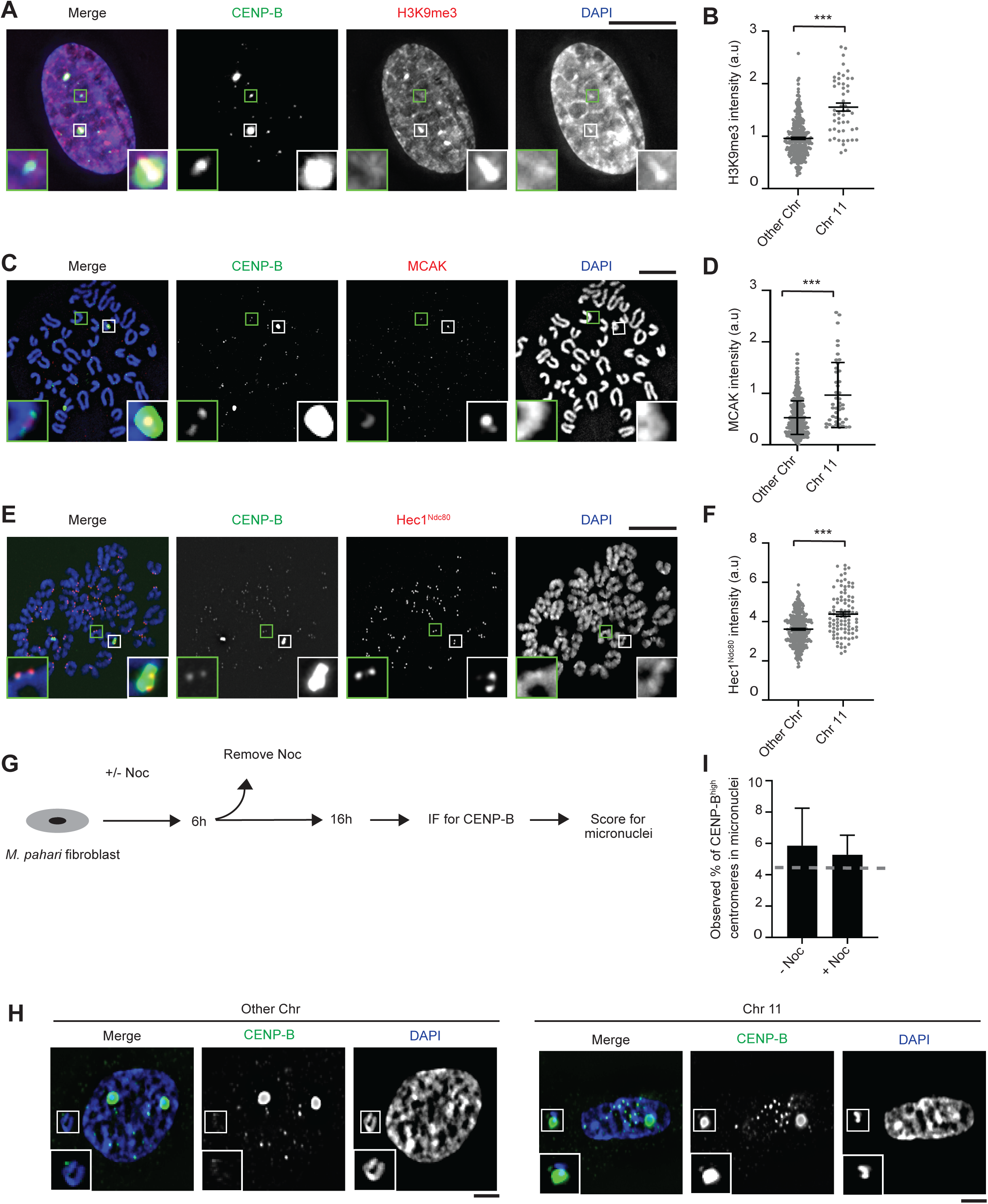
Chromosome 11 harbors levels of both pro and anti-microtubule binding proteins that are higher than on the other *M. pahari* centromeres. A) Immunofluorescence of H3K9Me3 from lung fibroblast cells derived from *M. pahari*. Insets: 4.0x magnification, Bar, 10 μm. Chromosomes are abbreviated as Chr in this figure. B) Quantification of H3K9Me3 from the experiment in panel A. The mean ratio (± SEM) is shown. n= 314 for the centromeres with low abundance of CENP-B and n= 50 for the centromeres with high abundance of CENP-B, pooled from 2 independent experiments. C) Immunofluorescence of MCAK from lung fibroblast cells derived from *M. pahari.* Insets: 6.5x magnification, Bar, 10 μm. D) Quantification of MCAK from the experiment in panel C. The mean ratio (± SEM) is shown. n= 389 for the centromeres with low abundance of CENP-B and n= 45 for the centromeres with high abundance of CENP-B, pooled from 2 independent experiments. E) Immunofluorescence of Hec1^Ndc80^ from lung fibroblast cells derived from *M. pahari*. Insets: 5.1x magnification, Bar, 10 μm. F) Quantification of Hec1^Ndc80^ from the experiment in panel E. The mean ratio (± SEM) is shown. n= 324 for the centromeres with low abundance of CENP-B and n= 94 for the centromeres with high abundance of CENP-B, pooled from 3 independent experiments. G) Schematic for measuring micronuclei containing chromosome 11 or other chromosomes. H) Immunofluorescence of micronuclei with low and high abundance of CENP-B centromeres from lung fibroblast cells derived from *M. pahari*. Insets: 1.8x magnification, Bar, 10 μm. I) Quantification of micronuclei from the experiment in panel H. Welch’s t test showed no significant difference between the actual micronuclei frequency and the expected frequency if there is no bias. A grey line represents the expected frequency given no bias, n= 133 (-Noc) and n= 419 (+Noc), pooled from 4 independent experiments.

Differences in centromere repeats between different mouse strains and species also have downstream molecular consequences that direct changes in the abundance of factors involved in microtubule-attachment (i.e. microtubule-binding proteins of the kinetochore, such as Hec1^Ndc80^) or in microtubule-destabilization (i.e. the kinesin, MCAK, that uses its motor activity to disassemble kinetochore microtubules) (*4, 7, 8*). The *M. pahari* genome harbors chromosomes with divergent centromere architectures that must undergo mitosis in unison and therefore it presents a unique opportunity for investigating the regulation of microtuble dynamics at the kinetochore. One likely scenario we considered is that the molecular changes yield a similar balance of microtubule couplers (e.g. Hec1^Ndc80^) and destabilizers (e.g. MCAK), so that their ratio is similar enough to each align and segregate on the mitotic spindle with similar fidelity. Current models suggest that CENP-B recruits MCAK is thought to be via its role in enriching H3K9me3 chromatin. Per our expectation, we observed an approximately 1.8-fold enrichment of MCAK on chromosome 11 relative to the other *M. pahari* chromsomes during mitosis (Fig. 6C,D). Thus, the heterochromatin pathway governing centromere strength leads to greater accumulation of a primary microtubule-destabilizer on chromosome 11. We hypothesized that the kinetochore pathway stimulated by CENP-B would likely be impacted, as well. To measure this, we detected the kinetochore microtubule coupler Hec1^Ndc80^, and found it, too, is recruited on 1.2-fold higher levels on chromosome 11 than on other chromosomes (Fig. 6E,F).

Since there are both increased levels of MCAK and Hec1^Ndc80^ on chromosome 11, we predict that this chromosome will properly segregate at rates comparable to the other *M. pahari* chromosomes. In unperturbed cells, chromosome segregation errors lead to a small percentage (1.5 +/-0.14% in our experiment) of cells having micronuclei. This is increased to 4.2 +/-0.91% in our experiment by transient incubation with the microtubule poison, nocodazole. In both cases, chromosome 11 missegregation to micronuclei (Fig. 6G-I) is near the expected value if there is no bias simply based on chromosome number (Fig. 6I, dashed grey line). Note, the slightly higher than expected value is explained by a likely undercount of the other chromosomes that are present in micronuclei since their levels of CENP-B which is used to identify missegregated chromosomes are lower than on chromosome 11. Together, our cell-based measurements support the notion that that despite chromosome 11 having different centromere DNA, its segregation fidelity is the same relative to the remaining centromeres.

## Discussion

For rapid centromere evolution to occur, a new innovation within a species would have to initiate on a single chromosome. Some innovations will strengthen centromeres and spread to other chromosomes, eventually becoming the dominant form within a species. We have identified and characterized a new repeat, π-sat^B^, in *M. pahari* that exists as a homogenous 6 Mbp array that confers centromere function on chromosome 11. The chromosome 11 centromere is an outlier compared to centromeres in other species. In *M. musculus,* all centromeres have similar numbers of CENP-B boxes and even in *H. sapiens* where centromeric sequence diverge, the range of CENP-B box numbers varies only ∼10 fold between chromosomes (*36, 38*). On the other hand, chromosome 11 has 27-143 times more CENP-B boxes than presumably evolutionarily older centromeres in *M. pahari* that we sequenced and assembled. The chromosome 11 centromere directly recruits high levels of CENP-B that, in turn, generates a larger kinetochore (Fig. 6E,F,7A). If not counterbalanced by corresponding increase in microtubule-destabilization, such an innovation would likely lead to mitotic chromosome segregation errors (Fig. 7B). We predict that other evolutionary instances of centromere innovation would require similar or analogous counterbalancing of these opposing molecular pathways during mitotic chromosome segregation, since imbalances are known to drive errors in mitotic chromosome segregation (*40*). Invasion of stronger centromere sequences into other chromosomes is likely to lead to imbalances during female meiosis that would favor the biased segregation of the new centromeres into the egg. Such a model would put female meiosis as the driver of the rapid expansion of new, stronger centromere sequences through an entire genome. Testing this model with *M. pahari* will require the identification (and/or isolation) of strains or closely related species where interspecies crosses produce viable animals with functional oocytes [note that *M. pahari* does not productively mate with *M. musculus* or *M. spretus* (unpublished observations, M.A.La.)]. Our study also opens up the prospect that other experimentally tractable model systems exist where centromere innovation similarly initiates from one specific chromosome.

**Figure 7:**
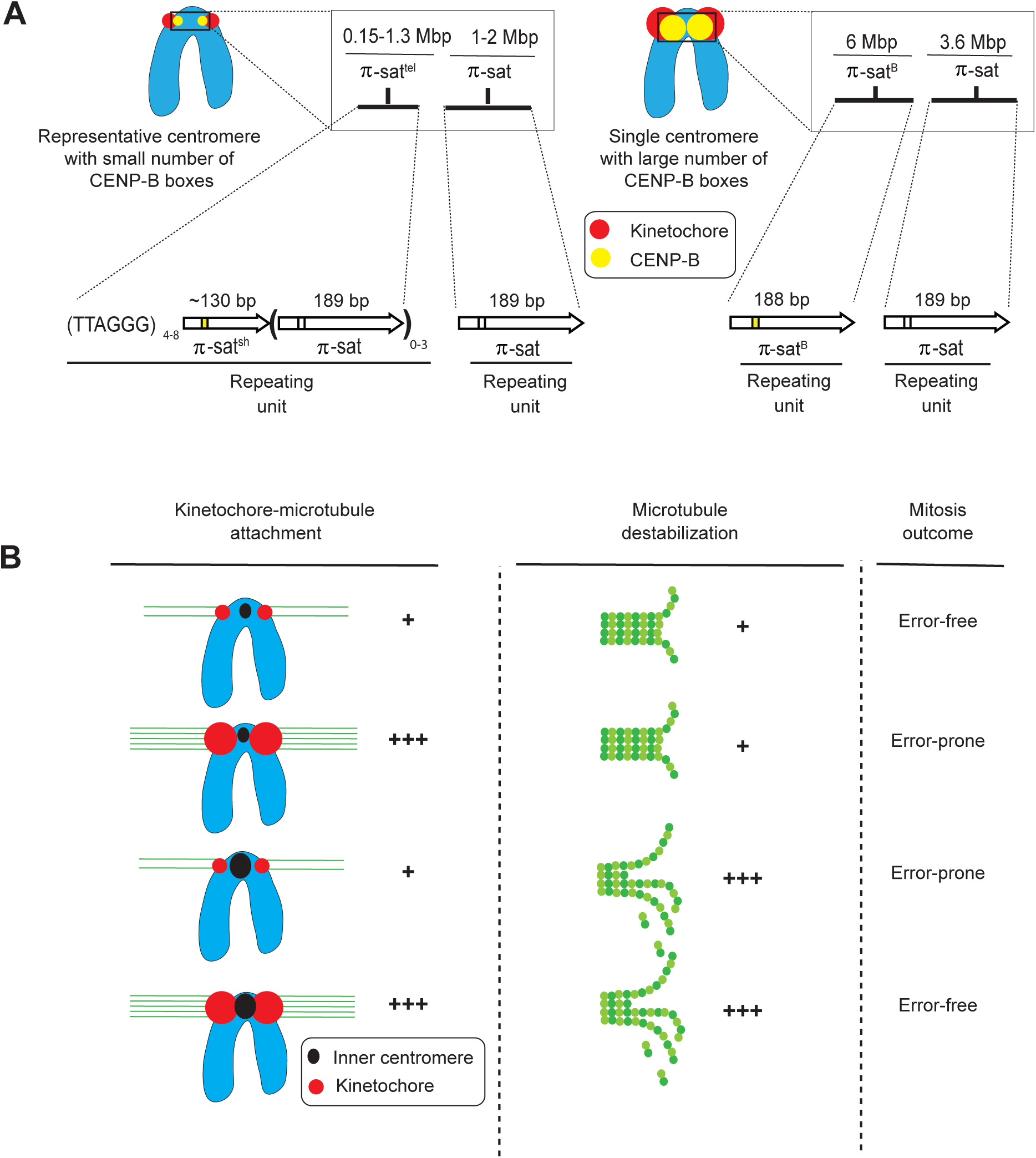
Divergent centromere DNA, molecular composition, and implications for mitotic chromosome segregation in *M. pahari*. A) Cartoon drawing summarizing the different types of *M. pahari* centromeres. The majority of *M. pahari* centromeres contain a low density of functional CENP-B boxes. Furthermore, these centromeres have two kinds of π-sat. First, the CENP-A containing region is a stretch of repeating units of π-sat that is short (∼130 bp) or long (189 bp) and interspersed with telomeric repeats. This is adjacent to a longer stretch of repeating units of 189 bp π-sat. The second type of *M. pahari* centromere has a high density of CENP-B boxes and is only found on chromosome 11. This centromere consist of 6 Mbp of homogenous π-sat^B^. The higher homogeneity of this centromeric DNA suggests that it is evolutionarily more recent relative to the other *M. pahari* centromeres. A) Model to understand different possible outcomes of centromere innovations during mitosis. The typical centromere has relatively low numbers of kinetochore attachments and relatively low amounts of microtubule destabilizer. These two factors balance each other allowing normal segregation during mitosis. If either pro- or anti-microtubule binding factors are increased in the absence of the other, there will be an imbalance resulting in incorrect segregation during mitosis. The chromosome 11 has higher levels of microtubule destabilizer and more microtubule attachments, but because both factors are increased together, the chromosomes can still undergo error-free mitosis.

To understand the arrangement of *M. pahari* centromeres, including the location of CENP-A nucleosomes, we started with information from short-read sequencing. In the final analysis, however, clarity on the situation would never have been achieved without employing long-read sequencing that yielded complete centromere assemblies. Our approach was modeled after the recent success in human centromere assemblies that has been a centerpiece accomplishment of the Telomere-to-Telomere consortium (*36–38*). Our work exemplifies how these approaches can be successfully employed to identify new centromere repeats in a non traditional model system (such as *M. pahari* that has had only modest genomic resources) for understanding mammalian chromosome evolution. Further, it succeeded in assembling centromeres harboring several megabases of repetitive DNA that are even more homogeneous in sequence than are human centromeres. For the older, more numerous *M. pahari* centromeres, our experiments revealed a association between CENP-A accumulation and repeats containing short spans of perfect telomere sequences as well as CENP-B boxes (Fig. 5). This suggests that for most centromeres in this species, the genetic contribution to centromere identity is particularly high. On the other hand, within the 6 Mbp of the most homogenous repeat, π-sat^B^ on chromosome 11, there is no strong sequence correlation with the specific peaks of CENP-A enrichment, since the sequences are almost identical at sites of either high or low CENP-A enrichment (Fig. 4A). It should be noted, though, that on chromosome 11 the highest peak of CENP-A enrichment is also adjacent to the telomere repeats at the natural telomere (Fig. 4). The lack of DNA sequence differences within the chromosome 11 centromere would suggest a strong epigenetic feedback that organizes the functional centromere at discrete sites within a large ‘sea’ of homogenized DNA repeats. Similar observations have been made in *M. spretus* (and *M. musculus*) where large stretches of homogenous centromeric DNA contain different kinds of chromatin at discrete locations despite no apparent sequence differences (*41*). On a technical note, our findings indicate that current sequencing methodologies and sequence assembly approaches can tackle some of the longest stretches of the most homogenized centromere sequences known in biology. Thus, massive stretches of similarly repetitive regions in other species (i.e. major satellite in *M. musculus*) should now be feasibly assembled using these methodologies.

Rapid centromere evolution is thought to be tied to karyotypic changes that separate closely related species (*42–46*). On one hand, the position (i.e. telocentric versus metacentric), size, and sequence of centromere DNA is malleable, since closely related species harbor striking changes between these attributes (*43, 46*). On the other hand, centromere repeats are generally homogenized within a species (*44*), supporting the concept that there is a positive functional consequence of having similar centromere function (i.e. recruitment of similar amounts of centromere proteins) across centromeres within a single species. The example in this study of *M. pahari*, shows how radical functional change to one of these attributes (centromere repeat) can be tolerated through counterbalancing pro-and anti-attachment of the centromere to spindle microtubules during cell division. We propose that the selective force to counterbalance functional centromere strength properties within a species shapes the nature and magnitude of innovations that would have the chance to ‘take hold’ in a population during the evolution of centromeres.

## Methods

### Experimental Model and Subject Details Mice

Mouse strains were purchased from the Jackson Laboratory (C57BL/6J # 000664 and PAHARI/EiJ # 002655). The CENP-B wild-type and knock-out mouse lines were generated as described previously (*20*). *M. pahari* used for ChIP were male and the age was 6 months. CENP-B WT/KO mice were female, and the age was 3.5 months. All animal experiments were approved by the Institutional Animal Care and Use Committee and were consistent with the National Institutes of Health guidelines.

### Cell lines

Primary lung fibroblasts were isolated from *M. musculus* or *M. pahari* as described previously (47). Cells were immortalized by transfection of SV40 large T antigen (48) a gift from Dr. B. Johnson (Upenn) using TransIT-X2 Dynamic Delivery System (Mirus). T-antigen integration was confirmed by PCR (5’-GGAATCTTTGCAGCTAATGGACCTTC-3’ and 5’ CCTCCAAAGTCAGGTTGATGAGCA-3’ primers yield a 246 bp product).

### Cell culture

The immortalized mouse primary fibroblasts were cultured in Dulbecco’s Modified Eagle medium (DMEM/F-12) supplemented with 10% FBS (Sigma), 1% penicillin–streptomycin (Gibco) at 37°C in a humidified atmosphere with 5% CO_2_.

### Primary mouse embryonic fibroblasts cultures

Mouse embryonic fibroblasts (MEFs) lines were isolated from a pregnant female E12.5-E13.5 embryos from *M. pahari* (PAHARI/EiJ). MEFs were cultured in MEF media composed of Dulbecco’s Modified Eagle medium (DMEM) supplemented with 10% FBS (Lonza), 100 μg/mL Primocin (Invivogen), and 1x GlutaMAX (Thermo Fisher Scientific/GIBCO) at 37°C in a humidified atmosphere with 5% CO_2_.

## Method Details

### MNase-digested chromatin and native ChIP

ChIP was performed as described previously (*4*). Briefly, nuclei were isolated from flash frozen mouse livers. Livers were homogenized in 4 mL ice-cold Buffer I (0.32 M sucrose, 60 mM KCl, 15 mM NaCl, 15 mM Tris-Cl, pH 7.5, 5 mM MgCl_2_, 0.1 mM EGTA, 0.5 mM DTT, 0.1 mM PMSF, 1 mM leupeptin/pepstatin, 1 mM aprotinin) per g of tissue by dounce homogenization. Homogenate was filtered through 100 μm cell strainer (Falcon) and centrifuged at 6000 × g for 10 min at 4°C. The pellet was resuspended in the same volume Buffer I. An equivalent volume ice-cold Buffer I supplemented with 0.2% IGEPAL was added, and samples were incubated on ice for 10 min. 4 mL nuclei were layered on top of 8 mL ice-cold Buffer III (1.2 M sucrose, 60 mM KCl, 15 mM NaCl, 5 mM MgCl_2_, 0.1 mM EGTA, 15 mM Tris, pH 7.5, 0.5 mM DTT, 0.1 mM PMSF, 1 mM leupeptin/pepstatin, 1 mM aprotinin) and centrifuged at 10,000 × g for 20 min at 4°C with no brake. Pelleted nuclei were resuspended in Buffer A (0.34 M sucrose, 15 mM HEPES, pH 7.4, 15 mM NaCl, 60 mM KCl, 4 mM MgCl_2_, 1 mM DTT, 0.1 mM PMSF, 1 mM leupeptin/pepstatin, 1 mM aprotinin), flash-frozen in liquid nitrogen, and stored at −80°C. Nuclei were digested with MNase (Affymetrix) using 0.05–0.15 U/μg chromatin in Buffer A supplemented with 3 mM CaCl_2_ for 10 min at 37°C. The reaction was quenched with 10 mM EGTA on ice for 5 min and an equal volume of 2× Post-MNase Buffer (40 mM Tris, pH 8.0, 220 mM NaCl, 4 mM EDTA, 2% Triton X-100, 0.5 mM DTT, 0.5 mM PMSF, 1 mM leupeptin/pepstatin, 1 mM aprotinin) was added prior to centrifugation at 18,800 × g for 15 min at 4°C. The supernatant containing the MNase-digested chromatin was pre-cleared with 100 μL 50% Protein G Sepharose bead (GE Healthcare) slurry in 1× Post-MNase Buffer for ∼ 2 hours at 4°C with rotation. Beads were blocked in NET Buffer (150 mM NaCl, 50 mM Tris, pH 7.5, 1 mM EDTA, 0.1% IGEPAL, 0.25% gelatin, and 0.03% NaN_3_). Pre cleared supernatant was divided so that an estimated 250 μg chromatin was used for ChIP 10 μg H3K9me3 antibody (Abcam ab8898) or 10 μg anti-mouse specific CENP-A antibody, (custom made by Covance and affinity-purified in-house) and 12.5 μg was saved as input. The custom polyclonal antibody raised against mouse CENP-A. Briefly, a New Zealand White rabbit was immunized using purified GST-tagged mouse CENP-A (aa 6-30) in PBS as an antigen and Freund’s adjuvant. ChIP samples were rotated at 4°C for 2 hours. Immunocomplexes were recovered by addition of 100 μL 50% NET-blocked protein G Sepharose bead slurry followed by overnight rotation at 4°C. The beads were washed three times with wash Buffer 1 (150 mM NaCl, 20 mM Tris-HCl, pH 8.0, 2 mM EDTA, 0.1% SDS, 1% Triton X-100), once with high salt Wash buffer (500 mM NaCl, 20 mM Tris-HCl, pH 8.0, 2 mM EDTA, 0.1% SDS, 1% Triton X-100), and the chromatin was eluted 2× each with 200 μL Elution Buffer (50 mM NaHCO_3_, 0.32 mM sucrose, 50 mM Tris, pH 8.0, 1 mM EDTA, 1% SDS) at 65°C for 10 min at 1500 rpm. The input sample was adjusted to a final volume of 400 μL with Elution Buffer. To each 400 μL input and ChIP sample, 16.8 μL of 5 M NaCl and 1 μLof RNAse A (10 mg/mL) was added. After 1 hour at 37°C, 4 μL of 0.5M EDTA and 12 μL Proteinase K (2.5 mg/mL, Roche) were added, and samples were incubated for another 2 hours at 42°C. The resulting Proteinase-K treated samples were subjected to a phenol-chloroform extraction followed by purification of DNA with a QiaQuick PCR Purification column (Qiagen) in preparation for high-throughput sequencing.

### High-throughput sequencing

Purified, unamplified input or ChIP DNA (see section MNase-digested chromatin and native ChIP) was quantified using an Agilent 2100 Bioanalyzer high sensitivity kit. DNA libraries were prepared for multiplexed sequencing according to Illumina recommendations as described (*49*) with minor modifications using NEB enzymes. Briefly, 5 ng input or ChIP DNA was end repaired and A-tailed. Illumina TruSeq adaptors were ligated, libraries were size-selected to exclude polynucleosomes, and adapter-modified DNA fragments were enriched by PCR using KAPA polymerase. Libraries were assessed by Bioanalyzer and the degree of nucleosome digestion for each experiment was assessed to avoid any potentially over-digested samples. Libraries were submitted for 150 bp, paired-end Illumina sequencing on a NextSeq 500 instrument.

### Paired-end sequencing analysis

Paired-end sequencing analysis was performed as described previously (4). Briefly, paired-end reads were converted to a name-sorted SAM file using picard-tools and samtools (50) then joined in MATLAB using the ‘localalign’ function to determine the overlapping region between the paired-end reads [requiring ≥95% overlap identity; (49)], and adapter sequences were removed if present. For analysis of minor and major satellite DNA, we used a custom tandem repeat analysis as described (49) with the following modifications. Joined reads were aligned to a trimerized mouse minor satellite consensus (GenBank: X14464.1) (30) or dimerized π-sat consensus or to the reverse complement of those tandem consensus sequences. Those joined reads aligning with ≥ 80% identity were chosen for further analysis. To calculate the percent of total reads, the number of joined reads aligning to the consensus sequence in either the forward or reverse complement orientation (without double-counting any joined read) was divided by the total number of joined reads. ChIP fold-enrichment was calculated as the fraction of reads mapping to minor satellite from the ChIP divided by the fraction of reads mapping to minor satellite in the input. Alignment of satellites was visualized with Matlab scripts Code3_plotting_fixIncrement_1sizeClass_JDM20170206_allPlots or 2020-04-29-INP-consensus align-hist-line. Logo’s were generated via Glam2 with the command (glam2 -2 -a 190 -b 220 n pahari_input_all_to_2nd_pisat_read.CENPBbox.10reads.fa -o 2nd_pisat_region_CBBox_10) (*51*). Sequence alignments were generated using CLC Sequence Viewer.

### TAREAN

Putative satellite sequences were identified with TAREAN (*35*) from Illumina input sequencing data (500,000 paired-end reads). Quality filtered and interlaced input fasta files were prepared from fastq files as recommended. TAREAN was run with the following parameters: cluster merging performed, no custom repeat database, cluster size threshold 0.0, no automatic filtering of abundant repeats, similarity search options: Illumina reads, read length 100 nt or more.

### ONT long-read sequencing of the M. pahari genome

To generate Oxford Nanopore Technologies (ONT) long-read sequencing data from the *M. pahari* genome, we first extracted high-molecular weight DNA from ∼2.5 million *M. pahari* liver nuclei by resuspending them in 1 mL of Puregene Cell Lysis Solution (Cat. # 158113) in a 2 mL microfuge tube. Then, we added 6 μL RNase A solution (Cat. # 158153) and incubated the mixture at 37°C for 40 min. We let the mixture cool to room temperature before adding 333 μL of Puregene Protein Precipitation Solution (Cat. # 158123), vortexing for 20 sec, and then placing the tube on ice for 10 min. We spun the tube containing the mixture at maximum speed in a 4°C microfuge for 3 min. Then, we split the supernatant into two separate 1.5 mL tubes with 700 μL in each. We added 750 μL isopropanol to each tube, inverted 50 times to mix, and then spun the tubes at maximum speed in a 4°C microfuge for 1 min. We discarded the supernatant and then added 666 μL 70% ethanol to one of the tubes. We vortexed the single tube for 1 second and then transferred all of the ethanol solution plus the pellet into the second tube. We vortexted the second tube for 1 second and then spun at maximum speed in a 4°C microfuge for 1 min. We washed the pelleted DNA with 666 μL 70% ethanol two more times (pouring off the supernatant, adding new 70% ethanol, briefly vortexing, and then spinning at maximum speed in a 4°C microfuge for 1 min). After the second wash, we removed as much ethanol as possible from the tube and let it air-dry for 25 min, until all traces of ethanol were gone. We then added 110 μL of Qiagen’s DNA Hydration Solution (Cat. # 158133) to the DNA pellet and stored it at 4°C for two days. Once the DNA was fully resuspended, we prepared the DNA for ONT long-read sequencing using the ONT ligation sequencing kit (Cat. # SQK-LSK109), following the manufacturer’s instructions. The library was loaded onto a primed FLO-MIN106 R9.4.1 flow cell for sequencing on the GridION. All ONT data was basecalled with Guppy 3.6.0 with the HAC model.

### PACBio HiFi Sequencing of the M. pahari genome

DNA extraction, library preparation, quality control, and sequencing were performed by the Genome Technologies Scientific Service at The Jackson Laboratory. Approximately 60 μg of high molecular weight DNA was isolated from spleen tissue of a single *M. pahari* (PAHARI/EiJ) male using the Monarch HMW DNA (NEB) according to the manufacturer’s protocols with 2000 rpm agitation speed. DNA concentration and quality were assessed using the Nanodrop 2000 spectrophotometer (Thermo Scientific; 434 ng/μL), the Qubit 3.0 dsDNA BR Assay (Thermo Scientific; 406 ng/μL), and the Genomic DNA ScreenTape Analysis Assay (Agilent Technologies). DNA quality was assessed to be high (260/280 = 1.83, 260/230 = 2.29) and suitable for input for PacBio HiFi library construction. A PacBio HiFi library was constructed using the SMRTbell Express Template Prep Kit 2.0 (Pacific Biosciences) according to the manufacturer’s protocols. Briefly, the protocol entails shearing DNA using the g-TUBE (Covaris), ligating PacBio specific barcoded adapters, and size selection on the Blue Pippin (Sage Science). The quality and concentration of the library were assessed using the Femto Pulse Genomic DNA 165 kb Kit (Agilent Technologies) and Qubit dsDNA HS Assay (ThermoFisher), respectively, according to the manufacturers’ instructions. The resultant library was sequenced on two SMRT cells on the Sequel II platform (Pacific Biosciences) using a 30 hours movie time. The two SMRT cells yielded 71.25 and 93.94 Gb of unique sequence data, respectively, with an average read length of 13.9 kb.

### Assembly of the M. pahari genome

We assembled the *M. pahari* genome using PacBio HiFi data and the whole-genome assembler, hifiasm [v0.16.1; (*39*)] using standard parameters. The assembled contigs were not scaffolded into the entire chromosomes.

### Alignment of CENP-A ChIP-seq and bulk nucleosomal data to the M. pahari genome assembly

To identify the location of centromeric chromatin, we took advantage of the *M. pahari* CENP-A ChIP-seq and bulk nucleosomal (input) data that we had generated. We first assessed the reads for quality using FastQC (https://github.com/s-andrews/FastQC), trimmed them with Sickle (https://github.com/najoshi/sickle) to remove low-quality 5’ and 3’ end bases, and trimmed them with Cutadapt (*52*) to remove adapters. We aligned the processed CENP-A ChIP-seq reads to the whole-genome *M. pahari* assembly using BWA (v0.7.17) with the following parameters: bwa mem -t {threads} -k 50 -c 1000000 {path_to_index} {path_to_read1.fastq} {path_to_read2.fastq}. We filtered the resulting SAM file to remove partial and supplementary alignments (retaining only primary alignments) with SAMtools flag-F4 before normalizing the data to the input data using DeepTools (v3.4.3) and the following command: bamCompare -b {path_to_CENP-A.bam} – b2 {path_to_input.bam} --operation ratio --binSize 5000 --minMappingQuality 60 -p 20 -o {out.bw}.

### Identifying CENP-B boxes and telomere repeats within the M. pahari sequence assembly

To identify the location of CENP-B boxes within the *M. pahari* genome assembly, we used a custom python script (findKmers.py) to detect the location of the following sequences within the assembly: 5’-TTCGNNNNANNCGGG-3’ (the 17-bp CENP-B box) and 5’-CCCGNNTNNNNCGAA-3’ (the reverse-complement of the 17-bp CENP-B box) or 5’-TTAGGG-3’ (telomere repeat) and 5’-CCCTAA-3’ (reverse complement of the telomere repeat). We ran the script with the following command: ./findKmers.py --kmers {CENP-B_box_sequences} --fasta {genome_assembly.fasta} -- out {out.bed}/. We visualized the resulting BED file on the UCSC genome browser with the *M. pahari* reference genome assembly.

### Metaphase chromosome spreads of MEFs, FISH, and image capture

FISH images of metaphase spreads of *M. pahari* cells were obtained using two different protocols. To obtain FISH images of π-sat, MEFs were cultured in MEF media to ∼ 80% confluency at 37°C in a humidified atmosphere with 5% CO_2_. Cells were subsequently serum starved on MEF media without FBS and exposed to 0.02 μg/mL Colcemid (Thermo Fisher Scientific/GIBCO) for 12 hours to synchronize and arrest cells in metaphase. MEFs were subsequently shaken off and resuspended in hypotonic solution (56 mM KCl) for 60 min. The harvested cells were then gradually fixed in 3:1 Methanol:Glacial Acetic Acid under constant agitation. Cells were pelleted by centrifugation, the fixative decanted off, and re-fixed for a total of 3-4 times. Following the final fixation round cells were suspended in 1-2 mL of fixative and dropped onto slides from a height of ∼1 m. Slides were allowed to air dry for approximately 10 min and then stored at -20°C until hybridization. Commercially synthesized oligos corresponding to the *M. pahari* sequence was PCR amplified and fluorescently labelled via nick translation. The Genomic DNA sequcence of putative *M. pahari* centromere sequence, π-sat, is; AAAACATGTATGTTTCTTCCTGCTCTATTAGACGCATTGTAAAGATATCTGTAGAACAAGCATAGGAATATGAGTGCACTTCTTGAAACACATGGTATTCTAAGAATAATTTCCTCCATGGCAGTTCAGAGCACTAAGTACAACTATGTGCACTCATGATTCACTCTGTTTTTCGTGAGTTTTGCATGT and the primers used were forward: 5’-AACATGTATGTTTCTTCCTGCTCT-3’, reverse: 5’-TGTACTTAGTGCTCTGAACTGCC-3’. Briefly, 250-1000 ng of PCR-amplified DNA was combined with nick translation buffer (200 mM Tris pH 7.5, 500 mM MgCl_2_, 5mM Dithiothreitol, and 500 mg/mL Bovine Serum Albumin), 0.2 mM dNTPs, 0.2 mM fluorescent nucleotides, 1 U DNAse (Promega), and 1 U DNA Pol I (Thermo Fisher Scientific). One of three fluorescent nucleotides was used for each satellite probe set: Fluorescein-12-dUTP (Thermo Fisher Scientific), ChromaTide Texas Red-12-dUTP (Thermo Fisher Scientific/Invitrogen), and Alexa Fluor 647-aha-dUTP (Thermo Fisher Scientific/Invitrogen). The reaction mixture was incubated at 14.5 °C for 90 min, and then terminated by addition of 10 mM EDTA. Probes ranged from 50-200 bp in size, as assessed by gel electrophoresis. Probes were used in FISH reactions on MEF metaphase cell spreads. Probes were denatured in hybridization buffer (50% formamide, 10% Dextran Sulfate, 2x saline-sodium citrate [SSC], and mouse Cot 1 DNA) at 72°C for 10 min and then allowed to re-anneal at 37°C until slides were ready for hybridization. Slides were dehydrated in a sequential ethanol series (70%, 90%, and 100%; each 5 min) and dried at 42°C. Slides were then denatured in 70% formamide/2x SSC at 72°C for 3 min, and immediately quenched in ice cold 70% ethanol for 5 min. Slides were subjected to a second ethanol dehydration series (90% and 100%; each 5 min) and air dried. The probe hybridization solution was then applied to the denatured slide. The hybridized region was then cover-slipped and sealed with rubber cement. Hybridization reactions were allowed to occur overnight in a humidified chamber at 37°C. After gently removing the rubber cement and soaking off coverslips, slides were washed 2 times in 50% formamide/2x SSC followed by an additional two washes in 2x SSC for 5 min at room temperature. Slides were counterstained in 80 ng/mL DAPI (Thermo Fisher Scientific/Invitrogen) for 10 min and air dried at room temperature. Lastly, slides were mounted with ProLong Gold AntiFade (Thermo Fisher Scientific/Invitrogen) and stored at -20°C until imaging. FISH reactions were imaged at 63x magnification on a Leica DM6B upright fluorescent microscope equipped with fluorescent filters (Leica model numbers: 11504203, 11504207, 11504164), LED illumination, and a cooled monochrome Leica DFC7000 GT 2.8 megapixel digital camera. Images were captured using LAS X (Version 3.7) at a resolution of 1920 x 1440 pixels.

FISH of π-sat^B^ and the π-sat^B^ CENP-B box was performed as described earlier (*53*) with some modifications. For FISH on metaphase spreads, *M. pahari* lung fibroblast cells were treated with 50 μM STLC (Sigma-Aldrich) for 2-4 hours to arrest cells during mitosis. Mitotic cells were blown off using a transfer pipette and swollen in a hypotonic buffer consisting of a of 75 mM KCl for 15 min. 3x10^4^ cells were cytospun in an EZ Single Cytofunnel in a Shandon Cytospin 4 onto an ethanol-washed positively charged glass slide and allowed to adhere for 1 min before permeabilizing with KCM buffer for 15 min. For interphase FISH, cells were seeded on a positvely charged glass slide before permabilizing with KCM buffer for 15 min. Slides were washed three times in KCM for 5 min at RT. Slides were fixed in 4% formaldehyde in PBS, before washing three times in dH_2_O for 1 min each. Slides were incubated with 5 μg/mL RNAse A in 2x SSC at 37°C for 5 min. Cells were subjected to an ethanol series to dehydrate the cells and then denatured in 70% formamide/2x SSC at 77°C for 2.5 min. Cells were dehydrated with an ethanol series.

Biotinylated π-sat^B^ DNA probe was generated by PCR using the template sequence TTTGAATCTAGATTTGTTTAGCTTAGAATACCATGTTTTCCGGAACTGCACTCATATTGATCTGCTTTTACTACAGAAATCTCTACAAAGCGTCTAATAGAGCAGGAAGAAAAATACCCGTTTTACACGAAAAACTCTTGAAATACAGAGTGAATCCTGAGTGCAGATACTTGTACTTAGTGCTCTGAACAAGAATTGAGGAATGTAAAGGATCCTAT and the primers used were forward: 5’-GTTTAGCTTAGAATACCATGTTT-3’ and reverse: 5’-TTCCTCAATTCTTGTTCAGAG-3’ with Biotin-11 dUTP (ThermoFischer Scientific; AM8450), purified with a G-50 spin column (Illustra), and ethanol-precipitated with salmon sperm DNA and Cot-1 DNA. Precipitated π-sat^B^ was suspended in 50% formamide/10% dextran sulfate in 2x SSC and denatured at 77°C for 5-10 min before being placed at 37 °C for at least 20 min. 100 ng DNA probe was incubated with the cells on a glass slide at 37°C overnight in a dark, humidified chamber. The CENP-B box probe was ordered from PNABio with a Cy3 fluorophore conjugated to the sequence TTTCGTGTAAAACGGGT. PNA probe was prepared as described previously (https://www.pnabio.com/pdf/FISH_protocol_PNABio.pdf). 50 μM of PNA probe was resuspended in formamide, heated to 55°C for 5 min and stored in aliquots at -80°C. After thawing, probe was diluted 1:100 in 10 mM Tris-HCl pH 7.2, 70% formamide, 10 mM maliec acid, 15 mM NaCl, and 0.5% blocking reagent (Roche 11096176001). Probe was denatured at 77°C for 5-10 min before being placed at 37°C for at least 20 min. 10 μL of probe was incubated with the cells on a glass slide at 37°C overnight in a dark, humidified chamber. The next day, slides were washed two times with 50% formamide in 2x SSC for 5 min at 45°C. Next, slides were washed two times with 0.1x SSC for 5 min at 45°C. Slides were blocked with 2.5% milk in 4x SSC with 0.1 Tween-20 for 10 min. For the π-sat^B^ FISH, Cells were incubated with NeutrAvidin-FITC (ThermoFisher Scientific; 31006) diluted to 25 μg/mL in 2.5% milk with 4x SSC and 0.1% Tween 20 for 1 hour at 37°C in a dark, humidified chamber. Cells were washed three times with 4x SSC and 0.1% Tween 20 at 45°C, DAPI-stained, and mounted on a glass coverslip with Vectashield (Vector Labs).

### Pulsed-Field gel electrophoresis and Southern blot

Pahari mouse genomic DNA was prepared in agarose plugs and digested with BstXI and HpaI enzymes by the manufacturer recommendation. The digested DNA was separated with the CHEF Mapper system (Bio-Rad; Run conditions for 5-1000 kbp range: 0.5x TBE, 1% pulse field certified agarose, 14°C, auto program, 16 hours run; Run conditions for 500-6000 kbp range: 1x TAE, 1% pulse field certified agarose, 14°C, 2 V/cm, 106° included angle, 5-40 min field switching with linear ramp, 92 hours run), transferred to a membrane (Amersham Hybond-N^+^), and blot hybridized with a 30 bp probe specific to the *M. pahari* centromeres (5’-TTCGTGTAAAACGGGTATTTTTCTTCCTGC-3’). To label the probe, 5’ and 3’ adapters below primers were added. The probe was labeled with ^32^P by PCR-amplifying a synthetic DNA template (5’-TTTGTGGAAGTGGACATTTCTTCGTGTAAAACGGGTATTTTTCTTCCTGCTAAAAATAGACAGAAGCATT-3’) with primers forward: 5’-TTTGTGGAAGTGGACATTTC-3’ and reverse: 5’-AATGCTTCTGTCTATTTTTA-3’. The blot was incubated for 2 hours at 65°C for pre-hybridization in Church’s buffer (0.5 M Na-phosphate buffer containing 7% SDS and 100 µg/mL of unlabeled salmon sperm carrier DNA). The labeled probe was heat denatured in a boiling water bath for 5 min and snap-cooled on ice. The probe was added to the hybridization Church’s buffer and allowed to hybridize for 48 hours at 65°C. The blot was washed twice in 2× SSC (300 mM NaCl, 30 mM sodium citrate, pH 7.0), 0.05% SDS for 10 min at room temperature, and four times in 2× SSC, 0.05% SDS for 5 min each at 60°C. The blot was exposed to X-ray film for 1-16 hours at – 80°C.

### Immunofluroscence and microscopy for immortalized mouse lung fibroblast cells

For a co-seed experiment involving CENP-A and CENP-B immunofluroscence, *M. pahari* and *M. musculus* immortalized lung fibroblast cells were co-plated in 1:1 ratio. For experiments involving H3K9Me3 immunofluroscence mouse lung fibroblast cells were fixed in 4% formaldehyde for 10 min at room temperature and quenched with 100 mM Tris (pH 7.5) for 5 min, followed by permeabilization with 0.5% Triton X-100 for 5 min at room temperature. All coverslips were then blocked in PBS supplemented with 2% fetal bovine serum, 2% bovine serum albumin, and 0.1% Tween before antibody incubation. The following primary antibodies were used: mouse mAb anti-mouse CENP-B (1:200, Santa Cruz Laboratories, (F4) sc-376283), rabbit pAb anti-mouse CENP-A (1:500, 0.535 μg/mL; custom-made by Covance and affinity-purified in house), rabbit pAb anti-human H3K9Me3 (1:500, Abcam ChIP grade ab-8898), rabbit pAb anti human MCAK a gift from D. Compton (Dartmouth), and rabbit pAb anti-mouse Hec1^Ndc80^ antibody (*54*). Secondary antibodies conjugated to flurophores were used: FITC Goat anti-Mouse (1:200, Jackson ImmunoResearch Laboratories # 115-095-146), Cy3 Goat anti-Rabbit (1:200, Jackson ImmunoResearch Laboratories #111-165-144). Samples were stained with DAPI before mounting with VectaShield medium (Vector Laboratories). For metaphase chromosome spread, cells were treated with 50 µM STLC for 4 hours to arrest the cells during mitosis. Mitotic cells were blown off using a transfer pipette and swollen in a hypotonic buffer consisting of a 1:1:1 ratio of 75 mM KCl, 0.8% NaCitrate, 3 mM CaCl_2_, and 1.5 mM MgCl_2_ for 15 min at room temperature. 5 x 10^4^ cells were cytospun onto a ethanol washed Superfrost Plus glass slide at 1500 rpm for 5 min and allowed to adhere for 2 min before fixing with 4% formaldehyde. Cells were permeabilized with 0.5% Triton X-100 for 15 min at room temperature followed by immunostaining. Images were captured at room temperature on an inverted fluroscence microscope (DFC9000 GT; Leica) equipped with a charge-coupled device camera (ORCA AG; Hamamatsu Photonics) and a 100x, 1.4 NA oil immersion objective. Images were collected as 0.2 μm Z-sections using identical acquisition conditions and Z series were deconvolved using LAS-X software (Leica). The fluroscence intensity was measured from deconvolved and maximum-projected images by ImageJ using 8 x 8 for H3K9Me3, 1.3 x 1.3 for MCAK, and 2.4 x 2.4 pixel box for Hec1^Ndc80^ using CENP-B as a refrence channel. The local background intensity was substracted from the measured fluroscence intensity. A minimum of 300 centromeres with low abundance of CENP-B and a minimum of 40 centromeres with high abundance of CENP-B were counted from at least two independent experiments. The mean ratio ± SEM is reported. For micronuclei experiment, *M. pahari* cells were arrested with nocodazole for 6 hours and then released for 16 hours. Cells were fixed with 4% formaldehyde for 10 min at room temperature and immunofluroscence was performed as described above.

## Acknowledgements

We thank our UPenn colleagues G. Birchak and K. McCannell for discussion, and L. Chmátal for isolating the *M. pahari* primary lung fibroblast cells. We also thank B. Sullivan (Duke) for sharing the protocol for mouse CENP-A antibody production, B. Johnson (Upenn) for providing the SV40 large T antigen plasmid, and D. Compton (Dartmouth) for providing the human MCAK antibody.

## Author contributions

C.W.G., N.P., J.M.D.-M., U.P.A., M.A.Li., M.A.La., G.A.L., B.L.D., and B.E.B. designed experiments. C.W.G., N.P., J.M.D.-M., U.P.A., M.A.Li., and G.A.L. performed experiments and analyzed data. J.M., P.L., and M.A.La. provided animal reagents. C.W.G., N.P., and B.E.B. wrote the paper. All authors edited the manuscript. B.E.B. directed the research.

## Data and materials availability

All sequencing data will be made publicly available at the time of publication on GenBank (centromere sequence assemblies) and SRA (raw sequencing files from Illumina, PACBio HiFi, and ONT). All data has been uploaded to PRJNA966193. All other data needed to evaluate the conclusions in the paper are present in the paper. The materials used in this study are available from commercial sources or from the corresponding author on reasonable request.

## Supplementary Figure Legends

Figure S1: π-sat sequence is almost identical to the top hit identified by the *k*-mer strategy Alignment of the satellites derived from the *k*-mer and TAREAN approach.

Figure S2: Functional CENP-B box is found at an pair of homologues containing π-sat^B^. Representative image of *M. pahari* fibroblast cells labeled with CENP-B box and π-sat^B^ FISH probes. Insets: 4.6x magnification. Bar, 10 μm.

Figure S3: Functional CENP-B boxes found at chromosome 11 differ from functional CENP-B boxes found on π-sat^tel^ on other centromeres.

Alignment of functional CENP-B box from π-sat^B^ and π-sat^tel^

Figure S4: Three additional *M. pahari* centromeres, all containing similar overall organization.

A-C) The fraction of π-sat repeats containing a functional CENP-B box (NTTCGNNNNANNCGGGN) and the frequency of telomeric repeats (TTAGGG) are shown. CENP-A ChIP-seq reads were aligned to the assembly revealing that CENP-A is primarily present on π-sat^tel^. A pairwise sequence identity heat map indicates the degree of homogeneity in centromeric DNA.

