## Supplemental Figures for "Centromere Innovations Within a Mouse Species"

### Figure S1

```
   k-mer  CATGATTCAC TCTGTTTTTT CTACTGAGTT TTGTGT--AA AACAAGTGAA TTTTCTTAAA GATCTTATTA  68
  TAREAN  CATGATTCAC TCTGTTTTTC --A-TGAGTT TTGTGTGTAA AACAAGTGAA TTT-CTTAAA GATCT-ATTA  65
   k-mer  GACACATTG- AGAGATTTTT TTGTAGAACA AGCATATAAT CATGAGTTTA GTTCCTGAAA TACTGAGTTA 137
  TAREAN  GACACATTGT AGAGATTTTT --GTAGAACA AGCATATGAA TATGAGTTTA GTTCCTAAAA TACTG-GTTA 132
   k-mer  ATTCTATGAA AACTNTCCAC AATTCTT--T CAGAGCAAAT AAGTACAAC- ATCT-----  188
  TAREAN  -TTCTATGAA AACATTCCAC AATTCTTGTT CAGAGCAA-T AAGTACAACT ATCTGCATT  189
```

### Figure S2

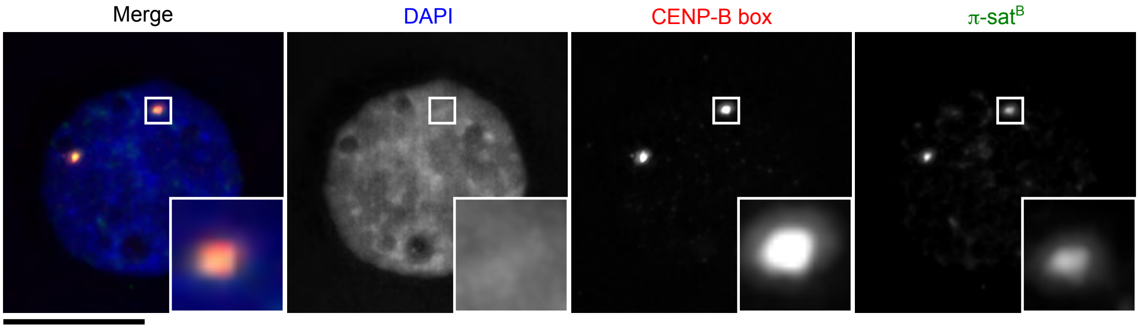

### Figure S3

A

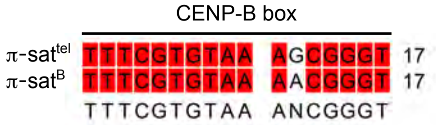
CENP-B box
π-sat^tel TTTCGTGTAA AGCGGGT 17
π-sat^B TTTCGTGTAA AACGGGT 17
TTTCGTGTAA ANCGGGT

### Figure S4

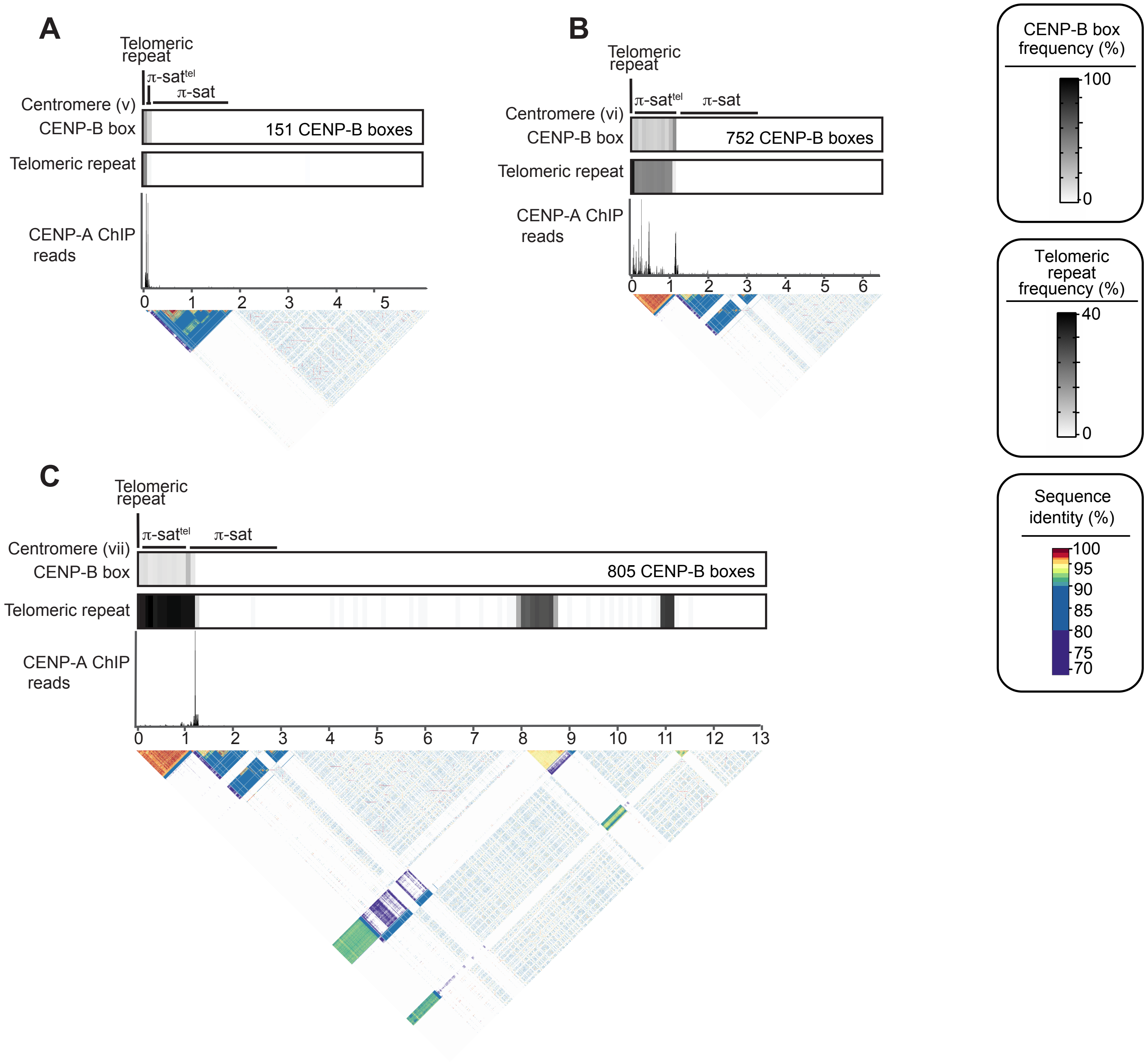

A
Telomeric repeat
π-sat^tel
π-sat
Centromere (v)
CENP-B box
151 CENP-B boxes
Telomeric repeat
CENP-A ChIP reads
0
1
2
3
4
5
B
Telomeric repeat
π-sat^tel
π-sat
Centromere (vi)
CENP-B box
752 CENP-B boxes
Telomeric repeat
CENP-A ChIP reads
0
1
2
3
4
5
6
C
Telomeric repeat
π-sat^tel
π-sat
Centromere (vii)
CENP-B box
805 CENP-B boxes
Telomeric repeat
CENP-A ChIP reads
0
1
2
3
4
5
6
7
8
9
10
11
12
13
CENP-B box frequency (%)
100
0
Telomeric repeat frequency (%)
40
0
Sequence identity (%)
100
95
90
85
80
75
70
